# Loss of Dnmt3a dependent methylation in inhibitory neurons impairs neural function through a mechanism that impacts Rett syndrome

**DOI:** 10.1101/815639

**Authors:** Laura A. Lavery, Kerstin Ure, Ying-Wooi Wan, Chongyuan Luo, Alexander J. Trostle, Wei Wang, Joanna Lopez, Jacinta Lucero, Mark A. Durham, Rosa Castanon, Joseph R. Nery, Zhandong Liu, Margaret A. Goodell, Joseph R. Ecker, M. Margarita Behrens, Huda Y. Zoghbi

## Abstract

Methylated cytosine is an effector of epigenetic gene regulation. In the mammalian brain, the DNA methyltransferase, Dnmt3a, is the sole “writer” of atypical non-CpG methylation (mCH), and methyl CpG binding protein 2 (MeCP2) is the only known “reader” for mCH. We set out to determine if MeCP2 is the sole reader for Dnmt3a dependent methylation by comparing mice lacking either Dnmt3a or MeCP2 in GABAergic inhibitory neurons. Loss of either the writer or the reader causes overlapping and distinct features from the behavioral to the molecular level. Loss of Dnmt3a results in global loss of mCH and a small subset of mCG sites resulting in more widespread transcriptional alterations and severe neurological dysfunction than seen upon MeCP2 loss. These data indicate that MeCP2 is responsible for reading part of the Dnmt3a dependent methylation in the brain. Importantly, the impact of MeCP2 on genes differentially expressed in both cKO models shows a strong dependence on mCH, but not Dnmt3a dependent mCG, consistent with mCH playing a central role in the pathogenesis of Rett Syndrome.

## Introduction

DNA methylation is a covalent epigenetic mark deposited mainly on the C5-position of naïve cytosine nucleotides in mammals. Although DNA methylation in mammals was described by Rollin Hotchkiss in 1948 (1), it took nearly thirty years to realize its function in the regulation of gene expression to shape fundamental biological processes including cell differentiation, genomic imprinting and genome stability (2, 3).

Today many features of the classic model for gene regulation by DNA methylation have proven more complex. Rather than static and solely repressive to gene expression, DNA methylation is functionally dynamic (4), and in some cases has been shown to activate gene expression (5–8) highlighting that context and protein factors that recognize methylated cytosine can alter the transcriptional outcome of this epigenetic mark.

It was also traditionally thought that DNA methylation in mammalian genomes could be found only at cytosines followed by guanine nucleotides (CpG methylation, or mCG), but it has become clear that cytosines in other contexts can also be methylated. This non-CpG methylation (also known as mCH, where H equals A, C or T) is not found in most mammalian cells but has been detected in pluripotent cells (9, 10), some human tissues (11), and is enriched in postnatal neurons (12–14). The mCH mark increases in neurons predominantly after birth and concurrent with the period of peak synaptogenesis throughout the first 25 years of life in humans (or first 4 weeks in mice). The resulting mCH pattern is largely conserved between mice and humans, where it makes up a high percentage of all DNA methylation in neurons (12,13,15). Although the timing and specification of mCH suggest it serves a critical function in maturing neurons, the molecular mechanism and functional consequences for loss of the mCH mark remain understudied.

In mammals, DNA methylation is catalyzed by *de novo* DNA methyltransferases Dnmt3a and Dnmt3b, and maintained by Dnmt1 in a CpG context (16). Recent studies have shown that Dnmt3a is the sole “writer” of mCH in postnatal neurons (13,17–19), where it is preferentially found in the CAC nucleotide context (12). Mutations in *DNMT3A* have recently been associated with disorders of neurodevelopment (20–22) suggesting that the mechanism by which Dnmt3a dependent methylation is written and read by other protein factors to regulate gene expression is critical for brain maturation. Loss of function of the only known “reader” of mCH in the mammalian brain, Methyl-CpG-binding protein 2 (MeCP2) (17,18,23), has long been associated with Rett syndrome (RTT) (24). RTT is an X-linked, postnatal neurological disorder; affected children are apparently healthy for the first 6-18 months of life, then lose their acquired milestones and develop a range of dysfunctions of the central and autonomic nervous systems (25, 26). Mouse models of *MECP2* mutations in female mice faithfully recapitulate patient symptoms, the severity of which, in humans and mice, is determined by the specific mutation and the pattern of X-inactivation (25,27,28). Numerous conditional knockout mice have revealed the function for MeCP2 in different cell populations in the brain as well as the etiology of RTT symptoms (29–41). Given the apparent need of every neural cell type for MeCP2, it is not surprising that *MECP2* mutations have been linked to other neuropsychiatric conditions beyond RTT (26, 28). Yet re-expression of MeCP2 in the CNS (42) rescues even the severe neurological symptoms and the prematurely lethal phenotype of constitutive null male mice (29, 41). RTT is thus driven by the loss of MeCP2 function in the nervous system, which continues to be essential into adulthood (43, 44), but the underlying architecture is sufficiently intact to support functional rescue.

Despite decades of research, the precise mechanism by which MeCP2 drives RTT phenotypes remains enigmatic. The recent finding that MeCP2 binds to mCH at disease-associated genes, and that mCH deposition occurs predominantly after birth and in conjunction with neuronal maturation, has led to the hypothesis that failure of mutant MeCP2 to bind to mCH could account for the postnatal onset of RTT symptoms (23). Whether loss of MeCP2 binding to mCH is sufficient to cause RTT has not been explored.

Here we set out to answer if MeCP2 is the only reader that interprets the unique methylation pattern set by Dnmt3a to direct gene expression in the maturing brain. If MeCP2 is the sole reader of these methylation marks, then loss of function of the writer should produce the same phenotype as loss of the reader at the behavioral and physiological level, as well as the same changes in gene expression at the molecular level. Such comparisons must be done in a cell-specific manner, since both mCH methylation (45) and the effects of MeCP2 loss are highly cell-specific (29–41). No one has performed a cell-type specific head-to-head comparison of knockout models at the behavioral, physiological and molecular level, which is necessary to define the functional relationship between Dnmt3a and MeCP2 in the nervous system. In this study, therefore, we systematically compared the effects of cell-specific knockout for *Dnmt3a* and *Mecp2* in mice of the same genetic background at the behavioral, physiological, and molecular level. We chose GABAergic inhibitory neurons for this comparison because knockout of *Mecp2* in this cell type recapitulates most of the RTT phenotype, including the delayed symptom onset (33), and loss of Dnmt3a and the methylation pattern it sets in inhibitory neurons has not been previously studied. Intriguingly, our findings indicate that while mCH has significant contribution to RTT, MeCP2 is responsible for enacting only a subset of Dnmt3a dependent gene regulation suggesting that there are other functional factors in this novel epigenetic pathway yet to be discovered that may impact neuropsychiatric phenotypes.

## Results

### Loss of Dnmt3a or MeCP2 in inhibitory neurons produces overlapping but not identical behavioral phenotypes

To compare the effects of loss of the mCH “writer” and “reader”, we deleted *Dnmt3a* or *Mecp2* utilizing floxed alleles for each gene in combination with a mouse line that drives Cre expression in GABAergic inhibitory neurons (*Viaat*-*Cre*) (29,33,46). We chose the *Viaat* promoter to drive Cre expression because it has been shown to turn on in the central nervous system around embryonic day 10 (47), before the earliest mCH marks have been detected in the brain (∼embryonic day 12.5 in the hindbrain) (15). We previously characterized mice with conditional knockout (cKO) of *Mecp2* in the same neurons on an F1 hybrid background (33), but to directly compare loss of the writer and reader we re-derived the *Mecp2* cKO on the C57BL/6J background. Conditional knockout mice showed loss of either Dnmt3a or MeCP2, respectively (Supplemental Figure 1A-B). We then tested both lines of mice for the same behavioral abnormalities previously reported (33).

Both cKO mice were born healthy but started to display symptoms around the time of weaning. Table 1 summarizes the phenotypes, showing that there is considerable overlap in the phenotypes but also specific features that are present in only one of the models (see Supplemental Table 1 for statistics). Both cKO mice first developed hind limb spasticity (Figure 1A), and by six weeks both *Dnmt3a* cKO and *Mecp2* cKO mice displayed obsessive grooming (Figure 1B), forepaw apraxia (Figure 1C), muscle weakness (Figure 1D), and motor deficits (Figure 1E, Supplemental Figure 2A). We found hippocampal-dependent, but not amygdala-dependent, learning and memory deficits (Figure 1F, Supplemental Figure 2C). Both mouse lines showed similar trends in anxiety-like behaviors (Supplemental Figure 2D-F), with *Mecp2* cKO achieving significance in one test (Supplemental Figure 2F). Starting at 7 to 8 weeks of age, the obsessive grooming led to skin lesions (Figure 1G); this self-injury was previously observed in *Mecp2* cKO mice on an F1 hybrid background (33) but was exacerbated on the C57BL/6J background, creating a need for humane euthanasia (see **Methods**). This shortened the period we were able to perform behavioral testing on both cKO mouse models. Both cKO mouse models still responded to pain stimuli (Supplemental Figure 2G-H), albeit to a lesser degree than controls. Rotarod and social interaction deficits that manifest in *Mecp2* cKO mice later in life on a F1 hybrid background did not develop within the foreshortened observation period (Supplemental Figure 2I-J).

**Figure 1.**
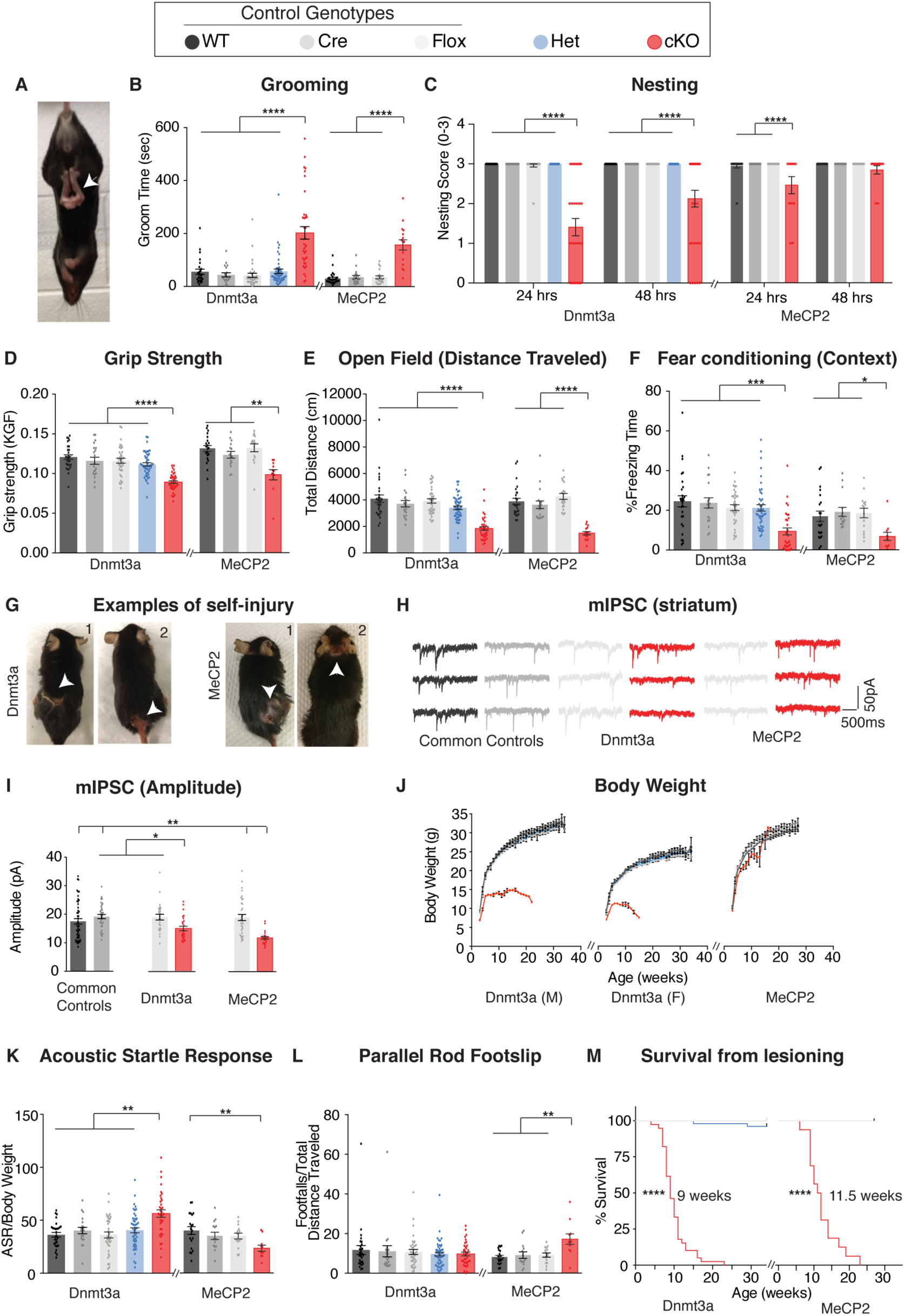
*Dnmt3a* cKO and *Mecp2* cKO mice show overlapping as well as distinct neurological deficits. **A)** Mice that lack Dnmt3a or MeCP2 in inhibitory neurons present with hindlimb spasticity. **B)** Obsessive grooming is increased in both knockout models. **C)** Nest building, **D)** grip strength, **E)** Open Field, **F)** Fear conditioning tests revealed impairments in both cKO lines. **G**) Self-injury in *Dnmt3a* cKO and *Mecp2* cKO mice necessitated humane euthanasia. **H)** Example traces of miniature inhibitory postsynaptic currents (mIPSCs) recorded in the dorsal striatum. Both cKO models show similar alterations in **I)** amplitude. **J)** Weekly body weight records for *Dnmt3a* cKO and *Mecp2* cKO mice showed only *Dnmt3a* cKO mice (here separated by sex-see **Methods**) were runted. **K**) *Dnmt3a* cKO and *Mecp2* cKO mice showed opposite alterations in acoustic startle response. **L)** Only *Mecp2* cKO mice displayed impairment on the parallel rod. **M)** *Dnmt3a* cKO mice had to undergo earlier euthanasia than *Mecp2* cKO mice due to the severity of their self-lesioning. *n= 11-52 (behavior), n=5-9mice per genotype with 24-50 neurons total (electrophysiology). *p<0.05, **p<0.01, ***p<0.001, ****p<0.0001. See Supplemental Table 1 for full statistics*.

**Table 1.** Summary of behavioral and physiological test results in cKO models.

| **Symptom Category** | **Test/observation** | **Result** |
| --- | --- | --- |
| General Health | Reduced body weight | *Dnmt3a* |
|  | Prematurely moribund (humane end due to lesioning) | both |
| Spasticity | Hind limb clasping | both |
| Repetitive Behavior | Forepaw stereotypies | *Dnmt3a* |
|  | Perseverative grooming & self-injury | both |
| Nociceptive pain | Delayed response to hot plate | both |
|  | Delayed response to tail flick | *Dnmt3a* |
| Apraxia | Poor nest building | both |
| Muscle Strength | Decreased grip strength | both |
| Motor | Rotarod | none |
|  | Parallel rod footslip- more footslips | *Mecp2* |
|  | Open field – decreased distance traveled | both |
|  | Open field – decreased vertical activity | both |
| Anxiety | Open field – time spent in center | none |
|  | Elevated plus maze- increased time in open arms | *Mecp2* |
|  | Light/dark box | none |
| Learning and Memory | Conditioned fear – decreased contextual fear response | both |
|  | Conditioned fear – cued | none |
|  | Conditioned fear – decreased response to tone #2 during fear learning (training) | *Dnmt3a* |
| Sensory processing | Increased acoustic startle response | *Dnmt3a* |
|  | Paired pulse inhibition | none |
| Social Behavior | Partition | none |
| Inhibitory signaling | Altered miniature inhibitory postsynaptic currents | both |

To compare cKO mouse models at the physiological level we measured miniature inhibitory postsynaptic currents (mIPSCs) in the dorsal striatum of *Dnmt3a* cKO, *Mecp2* cKO, and control mice. We observed decreased amplitude, as well as similar changes in frequency and charge (Figure 1H-I, Supplemental Figure 2K-L). Differences were not apparent in measurements of rise or decay time (Supplemental Figure 2M-N).

Despite these similarities, there were also clear differences between the cKO mice. *Dnmt3a* cKO mice were smaller than control mice starting at weaning age and throughout adulthood (Figure 1J). Forepaw stereotypies were evident in *Dnmt3a* cKO mice but not the *Mecp2* cKO mice on the C57BL/6J background (Supplemental Movie 1). Loss of Dnmt3a in inhibitory neurons led to an increased acoustic startle response (Figure 1K), with loss of MeCP2 in the same cell type showing the opposite trend, similar to a previous MeCP2 study (33). Only the *Mecp2* cKO mice developed motor incoordination (Figure 1L) and showed a trend for increased sensorimotor gating (Supplemental Figure 2O). *Dnmt3a* cKO mice also showed deficits in conditioned fear training (Supplemental Figure 2B). Finally, the *Dnmt3a* cKO mice showed much more severe self-injury, requiring humane euthanasia 2.5-week earlier than *Mecp2* cKO mice (Figure 1M).

### MeCP2 is a partial reader of Dnmt3a dependent methylation in striatal inhibitory neurons

Our behavioral data suggest that Dnmt3a and MeCP2 are functionally related, but it is still unclear what proportion of genes marked by Dnmt3a dependent methylation are regulated via MeCP2. To determine this, we tested our model that assumes MeCP2 is the only functional reader for these marks by sequencing the DNA methylome using a modified snmC-seq method adapted for bulk DNA samples (see **Methods**). We also performed RNA-seq on sorted GABAergic inhibitory neuronal nuclei from the striatum of *Dnmt3a* cKO, *Mecp2* cKO, and wild-type mice via the INTACT method (45) (Figure 2A). The striatum is ideal because mCH patterns are highly cell-specific and the vast majority of neurons (95%) in this region are inhibitory medium spiny neurons (48, 49). We found that methylation was stable in the absence of MeCP2 (Figure 2B, Supplemental Figure 3), but the Dnmt3a conditional knockout lost ∼90% of mCH (Figure 2C). We also noted a ∼10% loss of mCG in *Dnmt3a* cKO neurons (Figure 2D) consistent with partial mCG demethylation in the prenatal period followed by re-methylation later in development(15, 19). Dnmt3a is thus required to re-methylate mCG sites in the postnatal period. To determine if mCH and mCG marks written by Dnmt3a are coincident at genomic loci or independent of one another, we plotted the percentage of mCH versus mCG per gene in wild-type mice, as well as restricting the analysis to Dnmt3a dependent methylation (the change in methylation observed in the *Dnmt3a* cKO is defined as “Dnmt3a dependent”). While mCH and mCG in wild-type mice showed poor genome-wide correlation (Figure 2E), Dnmt3a dependent mCH and mCG showed significant correlation suggesting that the writing of mCH and mCG are coupled (Figure 2F).

**Figure 2.**
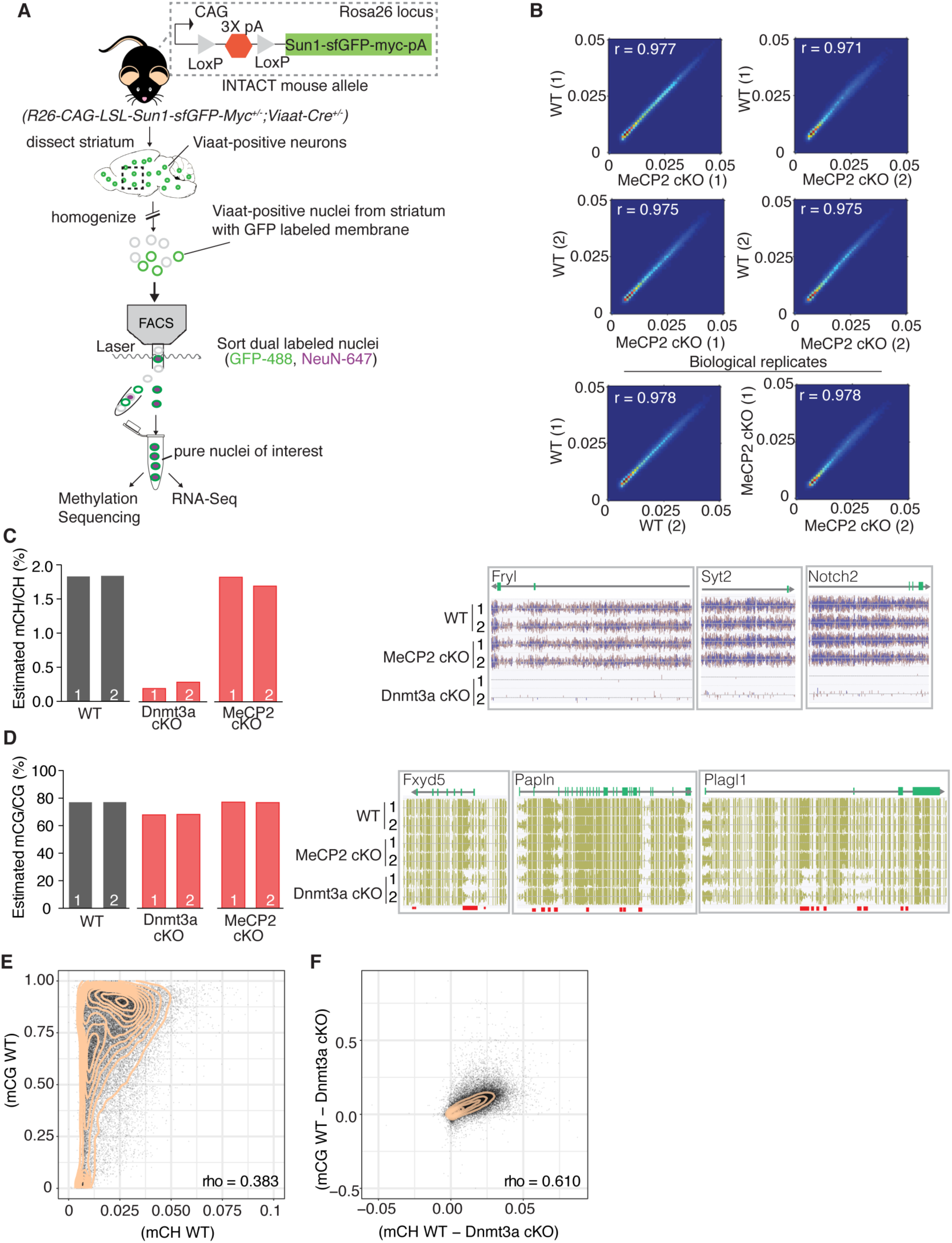
Methylation in striatal inhibitory neurons remains stable without MeCP2, but without Dnmt3a there is global loss of mCH and some loss of mCG. **A)** Schematic of INTACT method used to isolate inhibitory neurons from the striata of WT, *Dnmt3a* cKO, or *Mecp2* cKO mice that also conditionally express the INTACT allele. **B)** Spearman correlation of methylation profiles from inhibitory neurons sorted from WT vs. *Mecp2* cKO striatum, showing that methylation is stable in the absence of MeCP2. **C)** Bar graph showing the global mCH level in each biological replicate (left). Example genes from DNA methylome sequencing tracks showing mCH signal in two biological replicates per genotype (right). **D)** Bar graph showing the global mCG level in each biological replicate (left). Example genes from DNA methylome sequencing tracks showing mCG signal (right). Red bars indicate DMRs. **E)** Genome wide correlation between mCH and mCG in wild-type mice showing poor correlation. **F)** Genome wide correlation of Dnmt3a dependent mCG and mCG (the change in mCH methylation observed in the *Dnmt3a* cKO is defined as “Dnmt3a dependent”) showing good correlation to indicate that mCH and mCG written by Dnmt3a are coupled. Correlation values for E and F are Pearson correlations designated as “rho”. n= 2 mice per genotype (see methods for specific genotype information). See Supplemental Table 1 for replicate statistics.

Comparing gene expression by RNA-Seq for both knockout models, we found hundreds of differentially expressed genes (DEGs), either up- or down-regulated (Figure 3A, Supplemental Table 2). Although a large portion of DEGs in our *Mecp2* cKO model were significantly altered in *Dnmt3a* cKO mice (∼40%) indicating a dependence on Dnmt3a and an important contribution to RTT, we were surprised to find that only a small percentage of DEGs in our *Dnmt3a* cKO model were significantly altered in *Mecp2* cKO mice (i.e. are “MeCP2 dependent”, ∼12%) (Figure 3B). The amount of overlap was consistent at different p-value thresholds, and whether we considered only the DEGs that were up-regulated, down-regulated, or of certain length (Supplemental Figure 4A-C, Supplemental Table 2). Plotting the log2 fold-change in gene expression for each model against one another (Figure 3C), we found that DEGs common to both cKO models changed in the same direction and to a similar degree, consistent with an equal dependence on Dnmt3a and MeCP2. In all, our data show that MeCP2 is a restricted reader for Dnmt3a dependent methylation providing a platform for discovering novel functional factors in this pathway, as well as highlight a subset of genes regulated by MeCP2 that appear independent of Dnmt3a and the methylation it sets.

**Figure 3.**
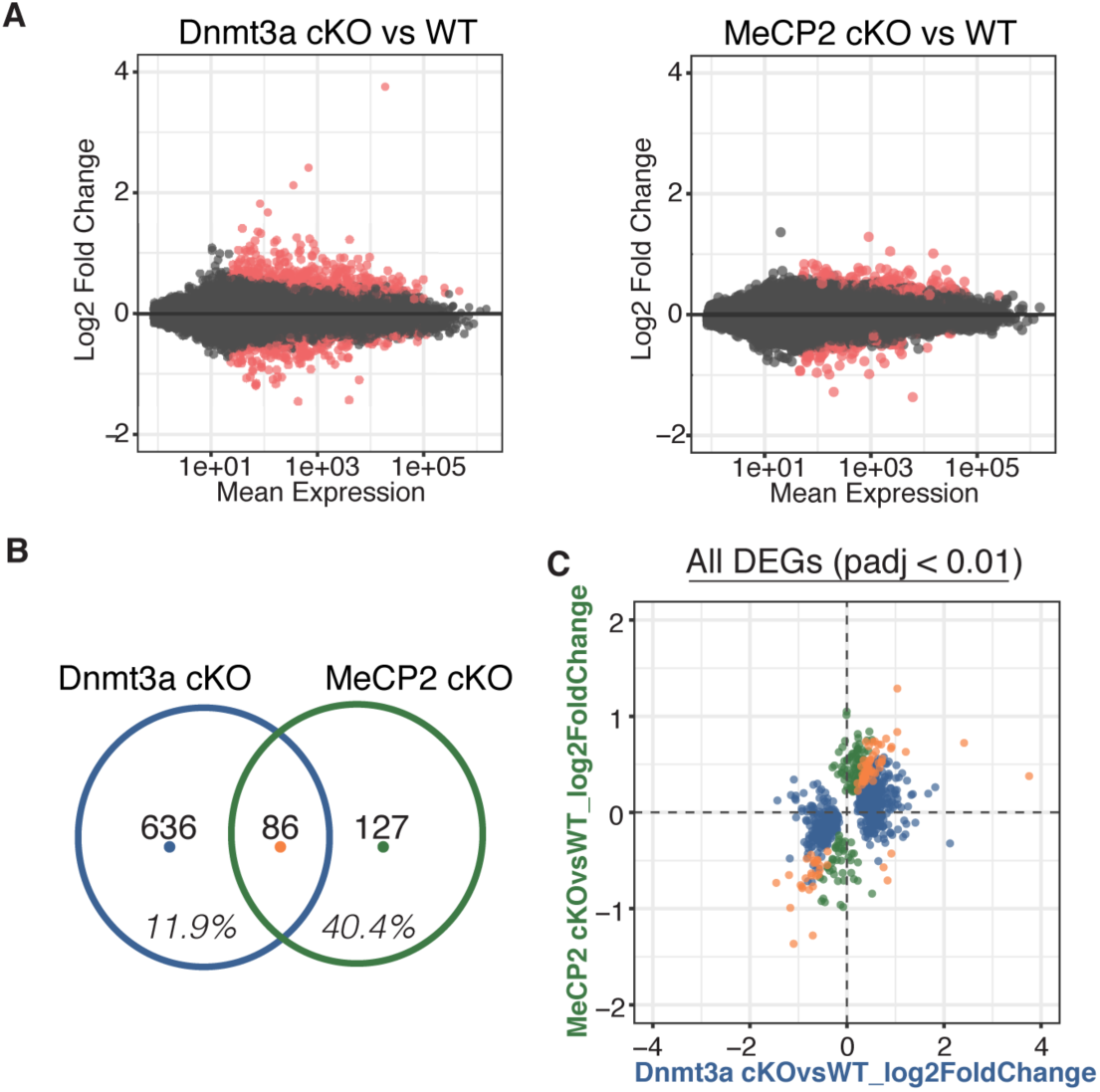
RNA-seq from sorted striatal inhibitory neurons of WT and cKO mice reveal MeCP2 is a restricted reader for Dnmt3a dependent methylation. **A)** RNA-seq data of sorted WT vs *Dnmt3a* cKO or *Mecp2* cKO striatal inhibitory neurons that also express the INTACT allele. Red dots represent genes with altered expression in the knockout cells (padj < 0.01). **B)** Differentially expressed genes (DEGs) that overlap between knockout models. Inhibitory neurons that lack MeCP2 or Dnmt3a share about 40% of the same DEGs. Only ∼12% of DEGs in inhibitory neurons that lack Dnmt3a are shared with neurons that lack MeCP2. **C)** Plot of log2 fold-change for DEGs in *Dnmt3a* cKO and *Mecp2* cKO models. DEGs that are only significantly misregulated in the *Dnmt3a* cKO model, only significantly misregulated in the *Mecp2* cKO model, or common to both models are colored in blue, green, or orange, respectively. The plot shows that the DEGs common to both models have similar degree and direction of change.

### Integrative gene expression and methylation analysis shows mCH and mCG loss contribute to Dnmt3a cKO DEGs, and reveals a strong mCH contribution to RTT

Plotting gene body mCH in wild-type mice for each category of differentially expressed genes, we find that, overall, DEGs have greater gene body mCH than genes that are unchanged, with up-regulated genes showing the biggest difference (Figure 4A). The same analysis showed that mCG levels are also higher on DEGs than non-DEGs for the majority of DEGs categories (Figure 4B), suggesting total methylation may also influence the final transcriptional outcome (50). Gene body methylation levels for all genotypes and statistics for comparisons can be found in Supplemental Figure 5. To elucidate the relative contribution of mCH or mCG written by Dnmt3a to gene expression changes in our cKO models, we employed a similar method to previous publications (19) and examined running average plots for the log2 fold-change in gene expression for each cKO model versus the percent change in Dnmt3a dependent methylation. These data were then fit with a univariate linear model to determine the percentage variance in log2 fold change explained by either mCH or mCG (R^2^, see **Methods**). For genes significantly misregulated only in the *Dnmt3a* cKO model we found a correlation with both Dnmt3a dependent mCH and mCG (Figure 4C), consistent with these marks being part of the same epigenetic program. These trends were statistically robust as when the same analysis was done 1000 times using random sets of non-DEG genes, the R^2^ values for our selected set of DEGs were significantly higher (Figure 4G, see **Methods**). The same trends hold when plotting the fold change in gene expression in the *Dnmt3a* cKO for genes that are commonly misregulated in both cKO models (Figure 4D,H). When we plot these commonly misregulated genes against the fold change in gene expression in the *MeCP2* cKO, we find a strong dependence on the change in gene expression to the change in mCH but not mCG (Figure 4E,I). As expected significant trends were not observed for Dnmt3a dependent methylation when plotting DEGs that were only significantly misregulated in the *MeCP2* cKO mice (Figure 4F,J). This integrative analysis demonstrates that loss of Dnmt3a dependent mCH and mCG significantly contributes to gene expression changes seen in the Dnmt3a cKO. Importantly we find that mCH, but not Dnmt3a dependent mCG, does in fact play a central role in mediating RTT pathogenesis.

**Figure 4.**
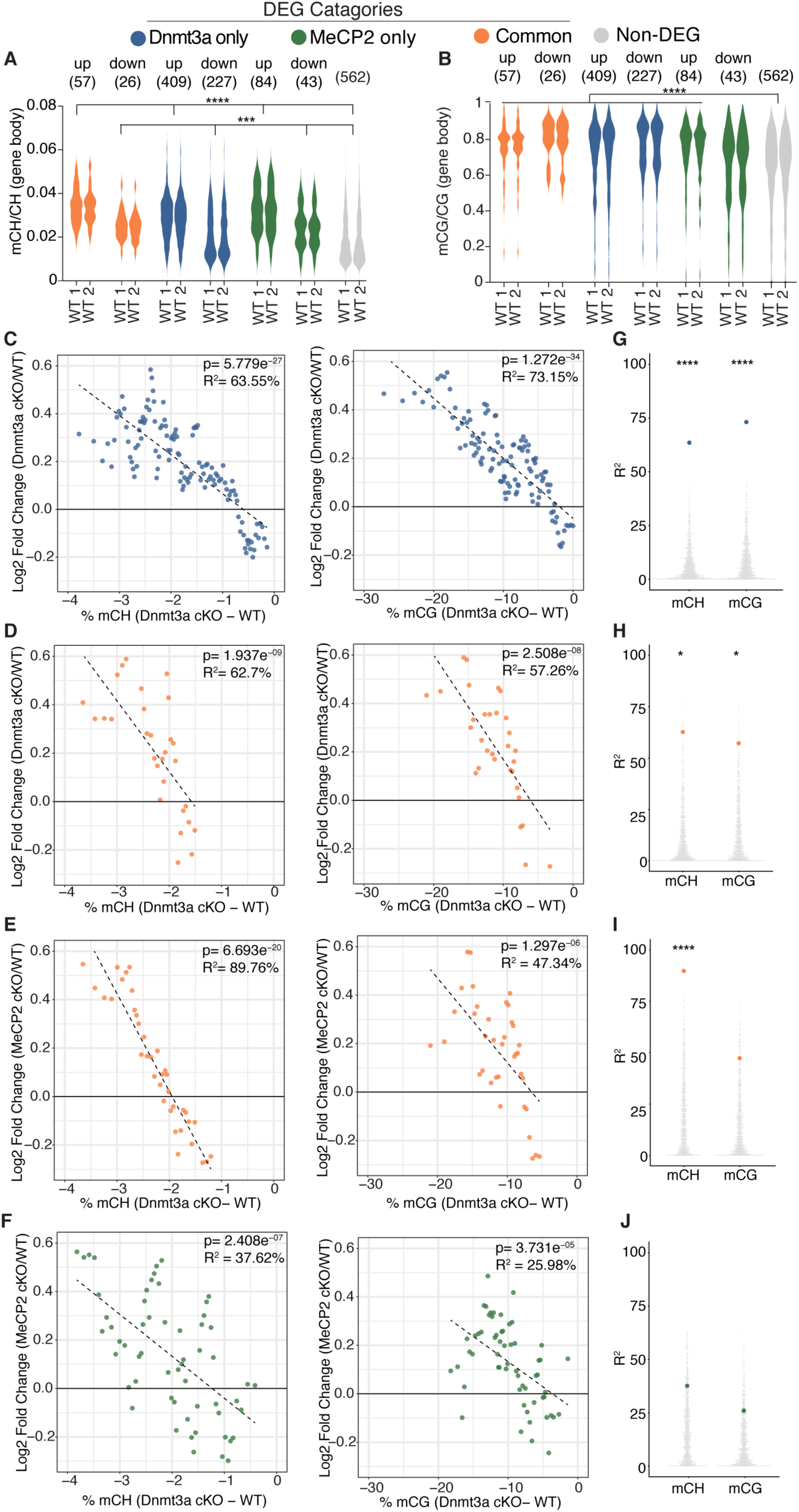
Integrative gene expression and methylation analysis shows mCH and mCG loss contribute to Dnmt3a cKO DEGs, and reveals a strong mCH contribution to RTT. **A)** Gene-body mCH levels in different categories of DEGs in WT mice are plotted, demonstrating that misregulated genes have higher mCH than genes that are unchanged. **B)** Gene-body mCG levels in different categories of DEGs in WT mice are plotted, demonstrating that misregulated genes (with the exception of MeCP2 down-regulated genes) have higher mCG than genes that are unchanged. **C)** Running average plot of log2fold change in gene expression for genes significantly misregulated only in the *Dnmt3a* cKO model versus the change in mCH methylation observed in the *Dnmt3a* cKO model (“Dnmt3a dependent mCH methylation”) (left). Running average plot of log2fold change in gene expression for these same genes versus the change in mCG methylation observed in the Dnmt3a cKO (“Dnmt3a dependent mCG methylation”) (right). **D)** Running average plots of log2fold change in gene expression in the *Dnmt3a* cKO model for genes commonly misregulated in both cKO models versus Dnmt3a dependent mCH (right) and mCG (left). **E)** Running average plots of log2fold change in gene expression in the *MeCP2* cKO for genes commonly misregulated in both cKO models versus Dnmt3a dependent mCH (right) and mCG (left). **F)** Running average plots of log2fold change in gene expression for genes that are only significantly misregulated in the *MeCP2* cKO model. **G-J)** R^2^ values from analysis in panels C-F shown as blue, orange or green dots, respectively, plotted over 1,000 random repetitions of the analysis with each repetition containing the same number of non-DEGs (padj>0.01). The results of random repetitions are shown as grey dots. *All plots were made with DEGs padj<0.01. n= 2 mice per genotype for methylation data. n= 4 mice per genotype (RNA-seq). *p<0.05, **p<0.01, ***p<0.001, ****p<0.0001*.

## Discussion

The discovery of mCH in postnatal neurons has shifted the view of how epigenetic gene regulation occurs in the brain. Beyond the obvious difference of unique context and cell-type restriction, the postnatal dynamics of this epigenetic program immediately suggest an important role for mCH in directing gene expression to ensure fully mature and functional neurons. Here we find that conditional knockout of the mCH writer, *Dnmt3a*, in GABAergic inhibitory neurons leads to genome-wide loss of mCH and a small subset of mCG sites, causing hundreds of gene expression changes and impairs neurophysiology and behavior with some deficits mirrored in a matched deletion of *Mecp2*. We were surprised to find only a modest contribution of MeCP2 to Dnmt3a/methylation dependent DEGs in our RNA-seq analysis. From this and our integrated gene expression and methylation analyses we distill two major conclusions. First, binding of MeCP2 to postnatal mCH contributes significantly to postnatal RTT symptoms, but there still appear to be other genomic targets recognized by MeCP2 or other functions whose loss contributes to the disease. Second, though MeCP2 “reads” a subset of Dnmt3a/methylation regulated genes, there appears to be a much broader gene regulatory program set up by the placement of global mCH and some mCG via Dnmt3a. These findings highlight the need to compare gene function at multiple biological scales, the molecular level in particular, when drawing conclusions about the functional relationship between proteins.

Uncovering the relationship between Dnmt3a and MeCP2 in the nervous system is challenging for a several reasons. First, while the phenotypes observed in whole-body knockout of *Mecp2* are recapitulated in the brain-specific knockout (29, 41), the phenotypes of whole-body (51) and brain-specific (52) knockout of *Dnmt3a* diverge, highlighting the essential role of Dnmt3a in peripheral tissues. In addition, while prior studies showed that early deletion of Dnmt3a in the brain drives neurological symptoms (52, 53), deletion of Dnmt3a using Cre-drivers that turn on postnatally have been demonstrated to produce minor (54) to no neurological symptoms (55). Therefore, it is likely that the underlying methylation pattern remains largely unaffected if Dnmt3a is deleted later in development, precluding the evolution of deficits. This is in contrast to MeCP2, where phenotypes are observed regardless of when the gene is deleted (43, 44). Our study deliberately eliminated these variations and shows that Dnmt3a and MeCP2 have overlapping but distinct roles to produce functionally normal neurons and behaviors. These data highlight the importance of extensive direct comparison between knockout models in a selected cell type, on the same genetic background, and to consider the developmental contribution for each gene being compared.

While DNA methylation remains stable in the absence of MeCP2, we find that loss of Dnmt3a in inhibitory neurons before birth leads to global loss of mCH, with a minor drop in mCG (∼10%), consistent with previous studies (13,15,19). The loss of mCG is likely due to de-methylation in the embryonic brain (13, 15) that cannot be re-methylated in the absence of Dnmt3a. At the RNA level, we show that loss of either Dnmt3a or MeCP2 leads to misregulation of hundreds of genes that normally have higher methylation (mCH in particular) relative to genes that are unchanged. We find expression of these genes changes in both the up and down direction in a manner that is independent of gene length. Importantly, the vast majority of the genes misregulated in both cKO models change to a similar degree and in the same direction, consistent with equal dependence of these genes on Dnmt3a and MeCP2 for proper regulation. While our data cannot rule out the possibility that secondary changes to gene expression may be occurring, the percentage of overlapping DEGs between cKO models is robust to p-value cutoff, direction of change or gene length. Integrating our genomic datasets, we find a significant contribution of mCH and mCG to the fold change in gene expression in the *Dnmt3a* cKO consistent with these marks functioning together to regulate gene expression during neuronal maturation. Intriguingly, we see a significant contribution of mCH, but not mCG to the misregulation of a subset of genes in the *MeCP2* cKO. This together with overlapping behavioral and physiological data suggests that binding of MeCP2 to mCH is critical for the postnatal evolution of RTT symptoms.

Our molecular data suggest that MeCP2 is likely one of a number of functional factors interpreting Dnmt3a dependent methylation. Though the overlap in gene expression changes between models has not been quantified until here, re-analysis of single-nuclear RNA-seq datasets from cortical inhibitory neurons (19) are consistent with our conclusion (Supplemental Figure 6). However, our data show a stronger dependence of MeCP2 DEGs on Dnmt3a function. With these data in hand we propose that the current model for Dnmt3a dependent gene regulation (19) be updated to incorporate our insights that MeCP2 is a restricted reader for Dnmt3a dependent gene regulation, as well as that postnatal mCG plays a role in the epigenetic program driven by Dnmt3a during neuron maturation.

There is a defined set of proteins in the mammalian genome that possess a methyl binding domain (MBD) and have been shown to bind mCH *in vitro* (56). These data highlight these proteins as primary candidates, though further studies are necessary to determine whether the other MBD-containing proteins read mCH in the brain, *in vivo*. However, there are other proteins involved in transcriptional regulation that do not possess a classic MBD and are sensitive to the methylation status of DNA, with some validated *in vivo* (57–61). For example, the DNA binding of many transcription factors is both positively and negatively affected by DNA methylation (58, 59), a phenomena that is not exclusive to mammals (62). Further, the DNA methylation and histone modification machinery are intimately linked (61). This link has been specifically shown in the differentiation of neural precursor cells, where methylation written by Dnmt3a antagonizes polycomb complexes to modulate gene expression (60). Given these findings, it is possible that similar mechanisms involving non-canonical methylation readers could be at play in Dnmt3a dependent gene regulation. Therefore, it will be important to undertake unbiased approaches to elucidate the full set of protein factors that drive gene regulation in this novel epigenetic pathway. All in all, our discovery that MeCP2 reads only some Dnmt3a dependent methylation marks lays the groundwork to uncover novel factors responsible for postnatal gene regulation that may be critical in setting up a mature, healthy mammalian brain.

## Materials and Methods

### Animal husbandry and handling

Baylor College of Medicine Institutional Animal Care and Use Committee (IACUC, Protocol AN-1013) approved all mouse care and manipulation. Mice were housed in an AAALAS-certified Level 3 facility on a 14hr light-10hr dark cycle. To obtain *Dnmt3a* cKO and control mice for behavior and electrophysiology, *Dnmt3a*^flox/flox^ mice(46) (C57BL6 background) were bred to *Viaat-Cre*^+/+^ mice(33) (C57BL/6J background) and *Dnmt3a*^flox/flox^ mice were bred to wild-type mice (C57BL/6J background) to get F1 *Dnmt3a*^flox/+^;*Viaat-Cre*^+/-^ females and *Dnmt3a*^flox/+^ males, respectively. F1 *Dnmt3a*^flox/+^;Viaat-Cre^+/-^females were mated with *Dnmt3a*^flox/+^ or *Dnmt3a*^flox/flox^ males to obtain F2 mice (males and females) of the following genotypes for behavior and electrophysiology: *Dnmt3a*^+/+^ (WT), *Dnmt3a*^+/+^;*Viaat-Cre*^+/-^ (Cre), *Dnmt3a*^flox/flox^ (Flox), *Dnmt3a*^flox/+^;*Viaat-Cre*^+/-^ (Het), *Dnmt3a*^flox/flox^;Viaat-Cre^+/-^ (cKO). Importantly, female mice carrying the Cre allele at F1 were used in F2 mating as we discovered that males of the same genotype (*Dnmt3a*^flox/+^;*Viaat-Cre*^+/-^) mated at F1 frequently transmit the Cre allele in the germline resulting in *Dnmt3a* null mice. To obtain *Mecp2* cKO and control mice behavior and electrophysiology, *Mecp2*^flox/flox^ females(29) (C57BL/6J background) were mated to *Viaat-Cre*^+/-^ males (C57BL/6J background) to get F1 male mice of the following genotypes: *Mecp2*^+/y^ (WT), *Mecp2*^+/y^;Viaat-Cre^+/-^ (Cre), *Mecp2*^flox/y^ (Flox), *Mecp2*^flox/y^;*Viaat-Cre*^+/-^ (cKO). Weekly weights and health scores were taken according to previously described disease score scale(63). For electrophysiological recordings control mice (WT and Cre) were shared for analysis and taken from both breeding schemes.

To obtain *Dnmt3a* cKO, *Mecp2* cKO and WT mice suitable for INTACT biochemistry experiments the breeding was as follows. *Dnmt3a*^flox/flox^ mice were bred to *R26-CAG-LSL-Sun1-sfGFP-Myc*^+/+^ mice(45) (C57BL/6J background) until both alleles were homozygosed to get *Dnmt3a*^flox/flox^;*R26-CAG-LSL-Sun1-sfGFP-Myc*^+/+^ mice. *Dnmt3a*^flox/flox^;*R26-CAG-LSL-Sun1-sfGFP-Myc*^+/+^ males were mated with *Dnmt3a*^flox/+^;*Viaat-Cre*^+/-^ females to obtain *Dnmt3a*^flox/flox^;*R26-CAG-LSL-Sun1-sfGFP-Myc*^+/-^;*Viaat-Cre*^+/-^ male mice used for biochemistry of striatal inhibitory neurons that lack Dnmt3a. *Mecp2*^flox/flox^ females were mated to *R26-CAG-LSL-Sun1-sfGFP-Myc*^+/+^ mice males until both alleles were homozygosed to obtain *Mecp2*^flox/flox^;*R26-CAG-LSL-Sun1-sfGFP-Myc*^+/+^ female mice. *Mecp2*^flox/flox^;*R26-CAG-LSL-Sun1- sfGFP-Myc*^+/+^ female mice were mated to *Viaat-Cre*^+/+^ males to get *Mecp2*^flox/y^;*R26-CAG-LSL-Sun1- sfGFP-Myc*^+/-^;*Viaat-Cre*^+/-^ male mice used for biochemistry of striatal inhibitory neurons that lack MeCP2. To obtain male mice used for biochemistry of wild-type striatal inhibitory neurons, *R26-CAG-LSL-Sun1-sfGFP-Myc*^+/+^ mice were bred to *Viaat-Cre*^+/+^ mice to get *R26-CAG-LSL-Sun1-sfGFP-Myc*^+/-^;*Viaat-Cre*^+/-^ male mice. For all cohorts of mice, mixed genotypes were housed separated by sex up to 6 mice per cage until 6-weeks of age and reduced to no more than 5 per cage after 6-weeks.

Before experimental cohorts were subject to investigation we noted *Dnmt3a* cKO and *Mecp2* cKO mice had a self-injury phenotype leading to skin lesions. Therefore, the institutional veterinarian was consulted to define humane euthanasia endpoint criteria for further experiments. Mice with lesions could be treated with antibiotic cream, and mice were euthanized when lesions extended below the skin layer. Survival age was taken as the age in which a mouse died of natural causes or when a mouse was euthanized due to skin lesions according to the institutional veterinarian guidelines set beforehand. Investigators were blind to genotypes of experimental mice until behavioral tests were finalized.

### Mouse Behavioral tests

*Dnmt3a* cKO, *Mecp2* cKO, and control mice were assigned individual ID numbers and the experimenters were blinded to genotype for the duration of behavioral testing. Experimental mice were divided into two cohorts for testing. Cohort 1 mice were subjected to the following tests, in order, at 6 weeks of age: open field, grooming, hotplate and tail flick, grip strength, parallel rod footslip, and prepulse inhibition/acoustic startle. The same mice were tested for conditioned fear at 8 weeks. Cohort 2 mice were subjected to the following tests, in order, at 6 weeks of age: elevated pus maze, light/dark box, rotarod, and partition/nesting. All behavioral assays were conducted during the light cycle, generally in the afternoon. All tests were conducted in light conditions.

All behavior data was analyzed using GraphPad Prism version 6 (GraphPad Software, La Jolla California USA, www.graphpad.com). Results were considered to be significant at p ˂ 0.05 and statistical significance reported in figures using a star notation to represent the lowest significance reached in the comparison (*\*p<0.05, **p<0.01, ***p<0.001, ****p<0.0001*). After behavioral analysis of the Dnmt3a cohort separated by both sex and genotype was completed, we determined that there was no statistical difference between males and females with the exception of body weight. Therefore, data for male and female mice were merged with the exception of body weight measures in the final analysis reported here. Statistical analysis for only males in the Dnmt3a cohort can also be found in Supplemental Table 1.

### Grooming

Mice were habituated in the test room for 30 minutes. Each mouse was individually placed in a clean housing cage without food, food grate, or water for 10 minutes. They were then videotaped for an additional 10 minutes. After recording, the mice were returned to their home cage. Videos were scored for grooming time by an investigator blind to the genotype of the test mouse. Data is shown as mean ± standard error of mean and was analyzed by one-way ANOVA with Tukey’s post hoc analysis.

### Grip Strength

Mice were habituated in the test room for 30 minutes. Each mouse was allowed to grab the bar of a digital grip strength meter (Columbus Instruments, Columbus, OH) with both forepaws while being held by the tail and then pulled horizontally away from the meter with a constant slow force until the forepaws released. The grip (in kg of force) was recorded and the procedure repeated for a total of three pulls. Data shown is the average of the three pulls presented as mean ± standard error of mean. Grip strength was analyzed by one-way ANOVA with Tukey’s post hoc analysis.

### Open Field

Mice were habituated for 30 minutes in the test room lit at 200 lux with white noise playing at 60 dB. Each mouse was placed singly in the open field apparatus (OmniTech Electronics, Columbus, OH) and allowed to move freely for 30 minutes. Locomotion parameters and zones were recorded using Fusion activity monitoring software. Data is shown as mean ± standard error of mean and was analyzed by one-way ANOVA with Tukey’s post hoc analysis.

### Fear Conditioning

Mice were habituated for 30 minutes outside the test room. Mice were placed singly into the conditioned fear apparatus (Coulbourn Instruments, Holliston, MA) that consisted of a lighted box with a floor made of parallel metal bars. On the training day, mice were placed in the chamber and subjected to two rounds of training, each of which consisted of 180 seconds of silence followed by a 30 second-long 80-85dB tone and 2 seconds of a 0.72 mA shock. 24 hours after training, the mice were returned to the box where they received the shock and freezing behavior recorded for six minutes. One hour later, the grated floor of the test chamber was covered, the shape changed with plastic panels, and vanilla scent added to the chamber. Mice were returned to the apparatus and subjected to a cue test consisting of 180 seconds of silence followed by 180 seconds of the original 80-85dB tone. Freezing behavior for all tests was scored using Freeze Frame 3 software (Actimetrics) with a threshold of 5.0. Data is shown as mean ± standard error of mean. Cue tests were analyzed by two-way ANOVA with Bonferroni’s post hoc analysis.

### Hot Plate and Tail Flick

Mice were habituated for 30 minutes in the test room prior to testing. Each mouse was placed individually on a 55°C hot plate (Stoelting Co., Wood Dale, IL) and observed for jumping, vocalization, hind paw lifting, or licking of the hind paws. At the first incidence of any of these behaviors the mouse was removed from the hot plate and the elapsed time noted. 30 minutes after the hot plate test, the mouse was placed on the tail flick apparatus (Stoelting Co., Wood Dale, IL) and restrained with a paper towel laid over the mouse and held down gently by the experimenter’s hand. The tail was laid in the groove above the lamp but was not restrained. The lamp was then turned on at 5-6eWatts; this also started the timer, which stopped automatically when the mouse moved its tail away from the lamp. The mouse was then returned to its home cage. Data was analyzed by one-way ANOVA with Tukey’s post hoc analysis.

### Elevated Plus Maze

Mice were habituated for 30 minutes in the test room lit at 200 lux with white noise playing at 60 dB. The elevated plus maze is a plus sign-shaped maze with two opposite arms enclosed by walls and two opposite arms open without walls. The entire maze is elevated above the floor. Mice were placed singly at the intersection of the four arms and allowed to move freely for 10 minutes. Activity was recorded by a suspended digital camera and recorded by the ANY-maze software (Stoelting Co., Wood Dale, IL). Data is shown as mean ± standard error of mean. Time and distance in the open arm were each analyzed by one-way ANOVA with Tukey’s post hoc analysis.

### Light/Dark Box

Mice were habituated for 30 minutes in the test room lit at 200 lux with white noise playing at 60 dB. Mice were placed singly in the light side of the light dark apparatus (Omnitech Electronics, Columbus, OH) and allowed to move freely for 10 minutes. Locomotion parameters and zones were recorded using Fusion activity monitoring software. Data is shown as mean ± standard error of mean. Time in Light was analyzed by one-way ANOVA with Tukey’s post hoc analysis.

### Parallel Rod Footslip

Mice were habituated in the test room for 30 minutes. Each mouse was placed in a footslip chamber consisting of a plexiglass box with a floor of parallel-positioned rods and allowed to move freely for 10 min. Movement was recorded by a suspended digital camera, while footslips were recorded using ANY-maze software (Stoelting Co.). At the completion of the test, mice were removed to their original home cage. Total footslips were normalized to the distance traveled for data analysis. Data is shown as mean ± standard error of mean and analyzed by one-way ANOVA with Tukey’s post hoc analysis.

### Acoustic Startle Response and Prepulse Inhibition

Mice were habituated for 30 minutes outside the test room. Each mouse was placed singly in SR-LAB PPI apparatus (San Diego Instruments, San Diego, CA), which consisted of a Plexiglass cylindrical tube in a sound-insulated lighted box. Once restrained in the tube, the test mouse was allowed to habituate for 5 minutes with 70 dB white noise playing. The mouse was presented with eight types of stimulus, each presented six times in pseudo-random order with a 10–20 ms inter-trial period: no sound, a 40 ms 120 db startle burst, three 20 ms prepulse sounds of 74, 78, and 82 dB each presented alone, and a combination of each of the three prepulse intensities presented 100 ms before the 120 dB startle burst. After the test, mice were returned to their home cage. The acoustic startle response was recorded every 1 ms during the 65 ms period following the onset of the startle stimulus and was calculated as the average response to the 120 db startle burst normalized to body weight. Percent prepulse inhibition was calculated using the following formula: (1-(averaged startle response to prepulse before startle stimulus/averaged response to startle stimulus)) x 100. Data are shown as mean ± standard error of mean. Percent prepulse inhibition was analyzed by two-way ANOVA with Bonferroni’s post hoc analysis, and acoustic startle response was analyzed by one-way ANOVA with Tukey’s post hoc analysis.

### Rotarod

Mice were placed on the rotating cylinder of an accelerating rotarod apparatus (Ugo Basile, Varese, Italy) and allowed to move freely as the rotation increased from 5 rpm to 40 rpm over a five-minute period. Latency to fall was recorded when the mouse fell from the rod or when the mouse had ridden the rotating rod for two revolutions without regaining control. This procedure was repeated for a total of four trials for two days. Data is shown as mean ± standard error of mean. Latency to fall was analyzed by two-way ANOVA with Bonferroni’s post hoc analysis.

### Partition test and nesting analysis

Mice were single-housed for 48 hours on one side of a standard housing cage. The cage was divided across its width by a divider with holes small enough to allow scent but no physical interaction. The test mouse was provided with a KimWipe folded in fourths as nesting material. At 24 hr and 48 hr of single-housing, the KimWipe was assessed for nesting score, as described previously(33). At least 16 hours before the partition test, a novel age- and gender-matched partner mouse of a different strain was placed on the opposite side of the partition. On the day of the test, the cage was placed on a well-lit flat surface. All nesting material, food pellets, and water bottles were removed from both sides of the cage, and the test mice were observed for 5 minutes while interaction time with the now-familiar partner mouse was recorded. Interactions involved the test mouse smelling, chewing, or actively exploring the partition. At the end of the first test (Familiar 1), a novel mouse of the same age, gender, and strain replaced the familiar partner mouse, and test mouse interactions were recorded for five minutes (Novel). The novel mouse was then removed and the familiar partner mouse returned to the cage, followed by observation for another 5 minutes (Familiar 2). At the completion of the partition test, test mice were returned to their original home cage. Data is shown as mean ± standard error of mean. Interaction times were analyzed by two-way ANOVA with Bonferroni’s post hoc analysis, and nesting scores were analyzed by one-way ANOVA with Tukey’s post hoc analysis.

### Western Blots

2-week old *Dnmt3a*^flox/flox^;*R26-CAG-LSL-Sun1-sfGFP-Myc*^+/-^;*Viaat-Cre*^+/-^ and *R26-CAG-LSL-Sun1- sfGFP-Myc*^+/-^;*Viaat-Cre*^+/-^ **OR** 6-week old *Mecp2*^flox/Y^;*R26-CAG-LSL-Sun1-sfGFP-Myc*^+/-^;*Viaat-Cre^+/-^*and *R26-CAG-LSL-Sun1-sfGFP-Myc*^+/-^;*Viaat-Cre*^+/-^ male were anaesthetize with isoflurane, whole brains dissected, split in half, frozen in liquid nitrogen and stored at −80°C till processing. Half-brains were thawed in 2mL lysis buffer (20mM HEPES pH 7.4, 200mM NaCl, 100mM Na_3_PO_4_, 10% Triton X-100, 1X protease inhibitor (Sigma Cat#5056489001), 1:2000 Universal Nuclease (Sigma Cat# 88701), dounced 50X with tight pestle and incubated on ice for 20min. Lysates were spun down 2x 20 minutes at 13,200 rpm at 4°C saving the supernatant each time. 20µg of protein was loaded onto a NuPAGE 4–12% Bis-Tris gradient gel and run in MES Running Buffer (Thermo Fisher Cat#NP0321BOX and NP000202, respectively). Protein bands were transferred to a nitrocellulose membrane in Tris-Glycine buffer plus10% methanol at 400mA for 1hr at 4°C. Membrane was blocked with 5% milk in tris buffered saline with 2% Tween-20 (TBST) for 1hr at room temperature (RT), cut and appropriate segments stained with primary antibodies overnight at 4°C. Membranes were washed in 1X TBST 3x for 5 minutes at RT and stained with secondary antibodies at 4°C for 1hr, followed by repeated washes in 1XTBST. ECL detection kit (Fisher 45010090) was used to detect HRP. Antibodies used were: 1:1000 anti-Dnmt3a (Santa Cruz 20703), 1:1000 anti-MeCP2 (in house N-terminal antibody), 1:20,000 anti-histone H3 (Abcam 1791) for primary antibodies, and 1:10,000 goat anti-rabbit-HRP (BioRad 170-5046) for the secondary antibody. Data is shown as mean ± standard error of mean and subject to unpaired t test with Welch’s correction for statistical analysis.

### Immunofluorescence

2-week old *Dnmt3a^f^*^lox/flox^;*R26-CAG-LSL-Sun1-sfGFP-Myc*^+/-^;*Viaat-Cre*^+/-^ and *R26-CAG-LSL-Sun1- sfGFP-Myc*^+/-^;*Viaat-Cre*^+/-^ **OR** 6-week old *Mecp2*^flox/Y^;*R26-CAG-LSL-Sun1-sfGFP-Myc*^+/-^;*Viaat-Cre*^+/-^ and *R26-CAG-LSL-Sun1-sfGFP-Myc*^+/-^;*Viaat-Cre*^+/-^ male were anaesthetize with Rodent Combo III and subject to transcardial perfusion using ice cold 1XPBS followed by PBS-buffered 4% paraformaldehyde, 10mL each. Brains were post-fixed in PBS-buffered 4% paraformaldehyde overnight at 4°C, equilibrated in 30% sucrose for 2 days and then frozen in Optimal Cutting Temperature medium (VWR 25608-930). For 2-week old brains, 25µm sagittal sections were taken on a Leica CM3050S cryostat, dried on slides and washed briefly in 1XPBS. Slides were transferred to antigen retrieval buffer (10mM Citrate pH6, 0.05% Tween-20) in a 95°C water bath for 20min, and then solution was allowed to cool to RT. For 6- week old brains 25µm sagittal sections were taken on a Leica CM3050S cryostat and then processed further as floating sections. All sections were blocked in blocking buffer (2% normal goat serum, 0.3% Triton X-100, in 1XPBS) for 1hr at RT and then stained with respective primary antibodies in blocking buffer (1:250 anti-Dnmt3a (VWR 64B1446) and 1:200 anti-Myc (Sigma C3956) for 2-week old brains and 1:500 anti-MeCP2 (Cell signaling 3456) and 1:800 anti-GFP (abcam 13970) for 6-week old brains). Sections were washed 4X in 1XTBST at RT for 5min each and then stained with secondary antibodies overnight at 4°C (1:1000 goat anti-mouse 555 (Invitrogen A21127) and 1:750 goat anti-rabbit 488 (Bethyl A120-201D2) for 2-week old brains and 1:750 goat anti-chicken 488 (Invitrogen A11039) and 1:1000 donkey anti Rabbit-568 (Invitrogen A10042) for 6-week old brains). Sections were washed 4x in 1XTBST at RT for 5min each and then stained with 2.5µg DAPI in 1XPBS then washed in 1XPBS, both for 10 minutes at RT. Sectioned were mounted in VECTASHIELD HardSet Antifade Mounting Medium without DAPI (Vector Laboratories H-1400) and let dry overnight. Images were taken on a Zeiss LSM 880 with Airyscan microscope at 63X. Images were taken with the same laser and gain settings and processed equivalently to facilitate comparison across genotypes. Of note, Dnmt3a related brains were processed at 2-weeks of age as Dnmt3a protein levels are at their peak, but then decline soon after this developmental time window(13) making Immunofluorescence challenging at 6-weeks of age.

### Electrophysiology

Acute fresh brain slices were prepared from 6-week old mice. Coronal slices (350 µm thick) containing striatum were cut with a vibratome (Leica Microsystems Inc., Buffalo Grove, IL) in a chamber filled with cutting solution containing 110mM C_5_H_14_ClNO, 25mM NaHCO_3_, 25mM D-glucose, 11.6mM C_6_H_7_O_6_Na, 7mM MgSO_4_, 3.1mM C_3_H_3_NaO_3_, 2.5 KCl, 1.25mM NaH_2_PO_4_ and 0.5mM CaCl_2_. The slices were then incubated in artificial cerebrospinal fluid (ACSF) containing 119mM NaCl, 26.2mM NaHCO_3_, 11mM D-glucose, 3mM KCl, 2mM CaCl_2_, 1mM MgSO_4_, 1.25mM NaH_2_PO_4_ at RT. The solutions were bubbled with 95% O_2_ and 5% CO_2_.

Whole-cell recording was made from medium spiny neurons in the dorsal striatum by using a patch-clamp amplifier (MultiClamp 700B, Molecular Devices, Union City, CA) under infrared differential interference contrast optics. Microelectrodes were made from borosilicate glass capillaries and had a resistance of 2.5-4 MΩ. Data was collected with a digitizer (DigiData 1440A, Molecular Devices). The analysis software pClamp10 (Molecular Devices) and Minianalysis 6.0.3 (Synaptosoft Inc., Decatur, GA) were used for data analysis. Miniature IPSCs were recorded in voltage-clamp mode in the presence of 10µM 6-cyano-7-nitroquinoxaline-2, 3-dione (CNQX), 50µM D-2-amino-5-phosphonopentanoic acid (AP5) and 1µM TTX. The glass pipettes were filled with high-Cl-intrapipette solution containing 145mM KCl, 10mM HEPES, 2mM MgCl2, 4mM MgATP, 0.3mM Na2GTP and 10mM Na2-phosphocreatine, pH 7.2 (with KOH). Signals were filtered at 2 KHz and sampled at 10 KHz. Data was discarded when the change in the series resistance was above 20% during the course of the experiment. The whole-cell recording was performed at 25 (±1) °C with the help of an automatic temperature controller (Warner Instruments, Hamden, CT). Data was analyzed with ordinary one-way ANOVA with Tukey’s multiple comparisons. Results were considered to be significant at p ˂ 0.05.

### Nuclei Isolation

Whole striatum was dissected from *Dnmt3a*^flox/flox^;*R26-CAG-LSL-Sun1-sfGFP-Myc*^+/-^;*Viaat-Cre*^+/-^, *Mecp2*^flox/y^;*R26-CAG-LSL-Sun1-sfGFP-Myc*^+/-^;*Viaat-Cre*^+/-^, and *R26-CAG-LSL-Sun1-sfGFP-Myc* ^+/-^;*Viaat-Cre*^+/-^, male mice in ice-cold HB buffer (0.25M sucrose, 25mM KCL, 5mM MgCl2, 20mM Tricine-NaOH) and flash frozen in liquid nitrogen and stored at −80°C. Experimental striatum (both halves of one mouse) were subjected to nuclear isolation and sorting for a total of 4 animals for each genotype. Individual striatum were dounced one at a time in 9mL lysis buffer (0.32M Sucrose, 5mM CaCl2, 3mM Mg(Ac)_2_, 0.1mM EDTA pH8, 10mM Tris-HCl pH8, 1mM DTT, 0.1% Triton X-100, 1X protease inhibitors (Sigma 5056489001), ribonuclease inhibitor (Promega N261A) 30 U/ml in DEPC treated water) in a 15mL dounce homogenizer (VWR 62400-642) 15 strokes with loose pestle, followed by 35 strokes with tight pestle. Homogenized tissue was gently layered onto two ultracentrifuge tubes (4.5 ml on each) filled with 8.5mL sucrose solution (1.8M sucrose, 3mM Mg(Ac)_2_, 1mM DTT, 10mM Tris-HCl pH8, 1X protease inhibitors (Sigma 5056489001), ribonuclease inhibitor 30 U/ml in DEPC treated water). Samples were spun in a Beckman Optima LE 80K ultracentrifuge for 2.5hrs at 6°C with slow deceleration. After centrifugation, the top layer of gradient and mitochondrial layer was discarded carefully with vacuum until ∼3mL remained. Remaining solution was gently poured off and tubes dabbed with a KimWipe being careful to not disturb the nuclear pellet. Nuclei were rehydrated on ice for 45 min in 500uL of rehydration buffer that was added drop-wise (0.5% BSA, 1X protease inhibitors, ribonuclease inhibitor 30 U/ml in 1X DPBS). Re-hydrated nuclei were pipetted up and down 50X with using a 1mL filter tip with the end cut to increase clearance size. Matched samples were then re-pooled and dual labeled with mouse anti-NeuN 647 (Millipore MAB377 labeled using Thermo Fisher Scientific A10475) and anti-GFP 488 (Thermo Fisher Scientific A-21311) at 1:300 and 1:1000 ratio, respectively, for 1hr at 4°C with gentle mixing. A BD Influx™ Cell Sorter at the Salk Flow Cytometry Core facility equipped with a 100 micron nozzle tip was used to isolate nuclei. Sheath fluid and pressure was 1XPBS (no Ca2^+^, Mg2^+^) and 18.5PSI, respectively. Nuclei were first gated based on light scatter properties to exclude debris (forward versus side scatter) then aggregate exclusion gating was applied (forward scatter as well as side scatter pulse width). Finally, nuclei were selected based on anti-NeuN647 and anti-GFP488 labeling. The nuclei fractions were collected at 4°C using a 1 or 2 drop purity sort mode and collected into rehydration buffer described above. Nuclei were then spun down at 5,000rpm for 15min at 4°C, solution removed until ∼50-100uL of buffer was left, and frozen on dry ice and stored at −80°C until DNA/RNA extraction for NGS library preparation.

### RNA extraction, NGS library preparation and RNA-seq

RNA was extracted from sorted nuclei using a single cell RNA purification kit from Norgen Biotek according to manufactures instructions and stored at −80C until library preparation at the Genomic and RNA Profiling Core at Baylor College of Medicine. The Genomic and RNA Profiling Core first conducted Sample Quality checks using the NanoDrop spectrophotometer and Agilent Bioanalyzer 2100. Total RNA was quanted by the user using Qubit 2.0 RNA quantitation assay. The NuGEN Ovation RNA-Seq v2 (protocol p/n 7102, kit p/n 7102-08) and the Rubicon ThruPlex DNA-Seq (protocol: QAM-108-002, kit p/n R400428) kits were used for library preparation as follows:

#### NuGEN Ovation RNA-Seq System v2 Protocol

Purified double-stranded cDNA was generated from approximately 5ng of total RNA and amplified using both oligo d(T) and random primers. Samples were quantified using the NanoDrop ND-2000 spectrophotometer and Qubit 2.0 DNA quantitation assay. One microgram of each sample’s ds-cDNA was sheared using the Covaris LE220 focused-ultrasonicator with a 400bp target size. The sheared samples were quantified using the Qubit 2.0 DNA quantitation assay. The fragment sizes were viewed on the Agilent Bioanalyzer to verify proper shearing.

#### Rubicon ThruPlex DNA-Seq Library Preparation Protocol

A double-stranded DNA library was generated from 50ng of sheared, double-stranded cDNA, preparing the fragments for hybridization onto a flowcell. This is achieved by first creating blunt ended fragments, then ligating stem-loop adapters with blocked 5’ ends to the 5’ end of the double-stranded cDNA, leaving a nick at the 3’ end. Finally, library synthesis extends the 3’ end of the double stranded cDNA and Illumina-compatible indexes are incorporated with 5 amplification cycles. The fragments are purified using AMPure XP Bead system. The resulting libraries are quantitated using the NanoDrop ND-1000 spectrophotometer and fragment size assessed with the Agilent Bioanalyzer. A qPCR quantitation is performed on the libraries to determine the concentration of adapter-ligated fragments using the Applied Biosystems ViiA7 TM Real-Time PCR System and a KAPA Library Quant Kit.

#### Cluster Generation by Bridge Amplification

Using the concentration from the ViiA7 qPCR machine above, 25pM from each equimolarly pooled library was loaded onto five lanes of a high output v4 flowcell (Illumina p/n PE-401-4001) and amplified by bridge amplification using the Illumina cBot machine (cBot protocol: PE_HiSeq_Cluster_Kit_v4_cBot_recipe_v9.0). PhiX Control v3 adapter-ligated library (Illumina p/n 15017666) is spiked-in at 2% by weight to ensure balanced diversity and to monitor clustering and sequencing performance. A paired-end 100 cycle run was used to sequence the flowcell on a HiSeq 2500 Sequencing System (Illumina p/n FC-401-4003).

### RNA-seq analysis

Adapter sequences were removed from raw sequencing reads using cutadapt (v1.13), trimmed reads were then aligned to reference genome GRCm38 (GENCODE vM15 Primary assembly) using STAR aligner(64) (version 2.5.3a) using default parameters. The number of reads aligned within the gene body (from TSS to TES) of each gene was tabulated using FeatureCount (65) (v1.5.3) (without extension on both ends). Finally, differential gene expression (DEG) analyses on the read counts were performed using DESeq2(66) (v1.6.2) in R environment. Genes with total read counts less 10 were filtered out from analysis. For analysis of differentially expressed genes split according to gene length as measured from transcription start site to transcription end site. Genes were defined as long or short according to length standards defined in Gabel et al(17).

### DNA Methylation Sequencing

DNA methylome libraries were generated using a modified snmC-seq method adapted for bulk DNA samples and as previous described in Sabbagh et al(67).1% unmethylated lambda DNA (Promega D1521) was added into each sample. Libraries were sequenced using an Illumina HiSeq 4000 instrument. The mapping of DNA methylome reads was done also described in Sabbagh et al(67).

Plots of mCH versus mCG in wild-type mice are made with gene body methylation values for percentage mCH or percentage mCG. All genes are plotted as one point per gene. Density contours are plotted with geom_density_2d in R and Pearson correlations were calculated with the cor function. The plots for the methylation differences are calculated with ((% mCH in WT) - (% mCH in cKO)) or as ((% mCG in WT) - (% mCG in cKO)).

### Integrative RNA-seq and methylation analysis

Methylation versus gene expression plots were made using running average binning on significantly differentially expressed genes (DEGs, padj < 0.01). To obtain bins, genes were first ordered based on their methylation value. Methylation is calculated as ((% mCH in WT) - (% mCH in cKO)) or as ((% mCG in WT) - (% mCG in cKO)). Genes were then binned such that each bin contained the same number of genes, with 80 percent overlap between consecutive bins. One point is plotted per bin. For the plots of genes significantly misregulated only in the *Dnmt3a* cKO model, each bin has 25 genes and the window moves by 5 genes per bin. For plots of genes significantly misregulated only in the *Mecp2* cKO model, as well as the common DEGs, the bin size is 10 genes and the window slides by 2 each time. The number of bins obtained for DEGs in the *Dnmt3a* cKO model were 121, and 42 for DEGs that were only significant in the *Dnmt3a* cKO model, and common DEGs, respectively. The number of bins obtained for DEGs in the *Mecp2* cKO model were 62, and 42 for DEGs that were only significant in the *Mecp2* cKO model, and common DEGs, respectively. After binning, a univariate linear model was fit to the data with the lm function in R, and the R^2^ (percentage variance in log2 fold change explained by methylation) was calculated.

To examine the significance of the observed trends in the running average plots, we compared their R^2^ values to R^2^ values from 1,000 random gene repetitions, with each repetition containing the same number of non-differentially expressed genes (padj > 0.01). Grey points represent each iteration of this process, with the original DEG (padj < 0.01) values for R^2^ shown in larger orange, green, or blue points. P value was then computed as (*r*+1)/(*n*+1) (68); r is number of repetitions where R^2^ is greater than that in the DEGs, and n is total number of repetitions.

## Acknowledgments

We are grateful to members of the Zoghbi lab for helpful discussions, Drs. H. K. Yalamanchili and H-H. Jeong for assistance with integrative analysis of RNA-seq and methylation data, and V.L. Brandt for critical comments on the manuscript. We thank Drs. Michael Greenberg and Hume Stroud for assistance with access to and discussion of published single-nuclei RNA-seq data. We thank the Neurovisualization and Neurobehavioral Cores at the Jan and Dan Duncan Neurological Research Institute at Texas Children’s Hospital and the BCM-IDDRC (U.S. NIH Grant U54HD083092). This project was supported by NIH/NINDS 5R01NS057819-13 (HYZ), NIH/ NIMH R01MH112763 (MMB and JRE), the Genomic and RNA Profiling Core at Baylor College of Medicine, and the Flow Cytometry Core Facility of the Salk Institute with funding from NIH-NCI CCSG: P30 014195. LAL is a Howard Hughes Medical Institute Fellow of the Life Sciences Research Foundation. HYZ and JRE are investigators with the Howard Hughes Medical Institute.

## Author Contributions

LAL and HYZ conceived of the study. LAL and KU collected and analyzed behavioral data. WW and JL collected and analyzed electrophysiology data. LAL and JL collected and processed samples for RNA and methylation sequencing. RC prepared and JRN sequenced methylation libraries. YW processed and analyzed RNA-seq data. CL processed and analyzed genomic methylation data. AT made mCH versus mCG plots and preformed integrative RNA-seq and methylation analysis. MAD assisted in tissue collection. LAL and HYZ wrote the manuscript. LAL, KU, YW, CL, WW, MAD, ZDL, MAG, JRE, MMB and HYZ interpreted data, edited manuscript and all authors approved of the final version.

## Data Access

DNA methylome data can be accessed through a web browser at http://neomorph.salk.edu/Striatum_Inhibitory_Neuron.php and at the Gene Expression Omnibus database (GEO) at accession number GSE124009.

RNA-seq data can be accessed at GEO at accession number GSE12394.

---

**Supplemental Figure 1.**
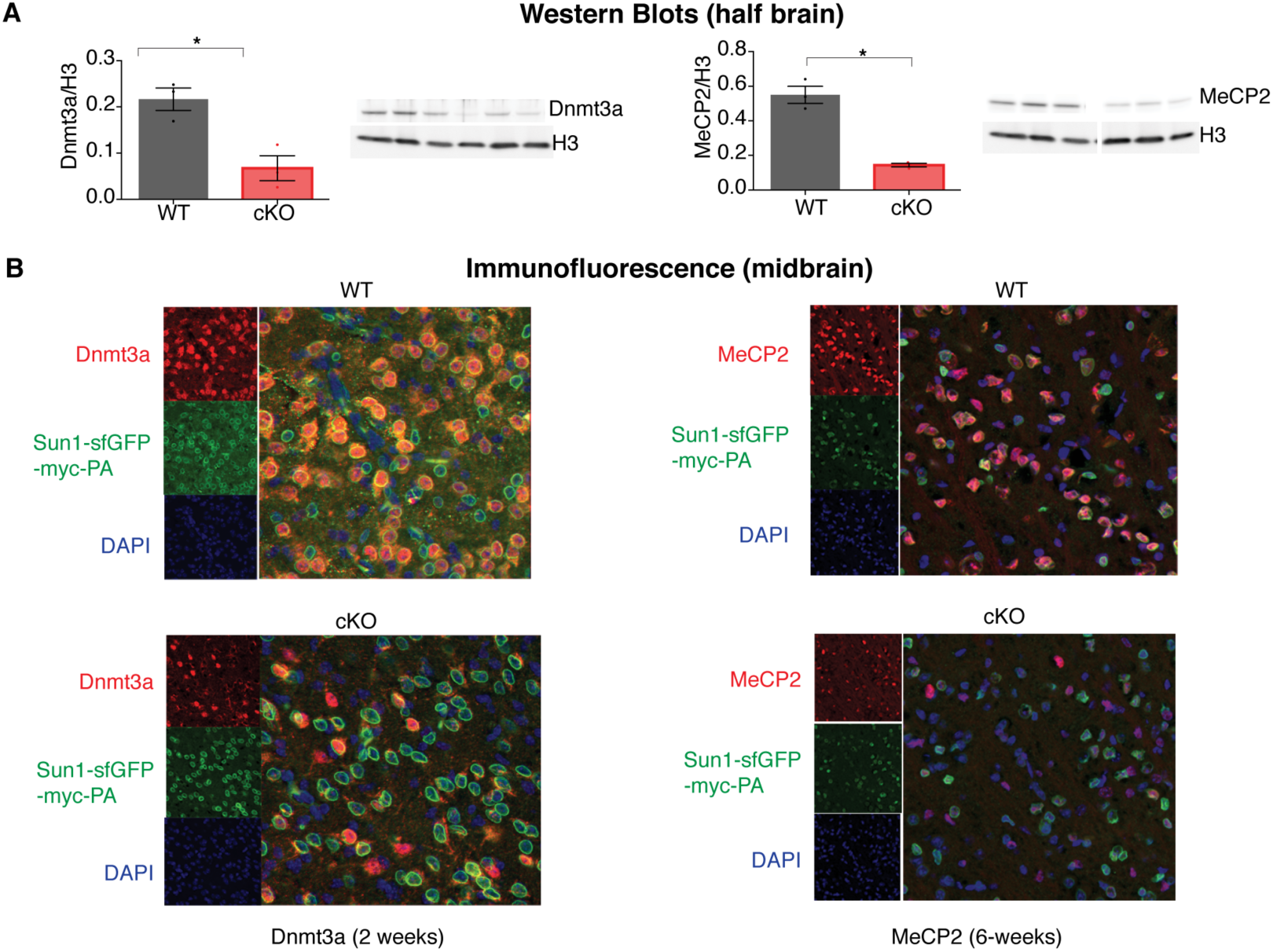
**A)** Western blot of half brain hemisphere from *Dnmt3a* or *Mecp2* cKO mice demonstrating appropriate loss of each protein. **B)** Immunofluorescence (IF) images of WT, *Dnmt3a* cKO or *Mecp2* cKO mice probing for Dnmt3a or MeCP2 (red), the Sun1-sfGFP-myc-PA fusion protein that marks the nuclear envelope and is dependent on Cre expression (green), and DAPI to mark genomic DNA (blue). n=3 mice per genotype (western), n=3 mice (IF) representative image shown.

**Supplemental Figure 2.**
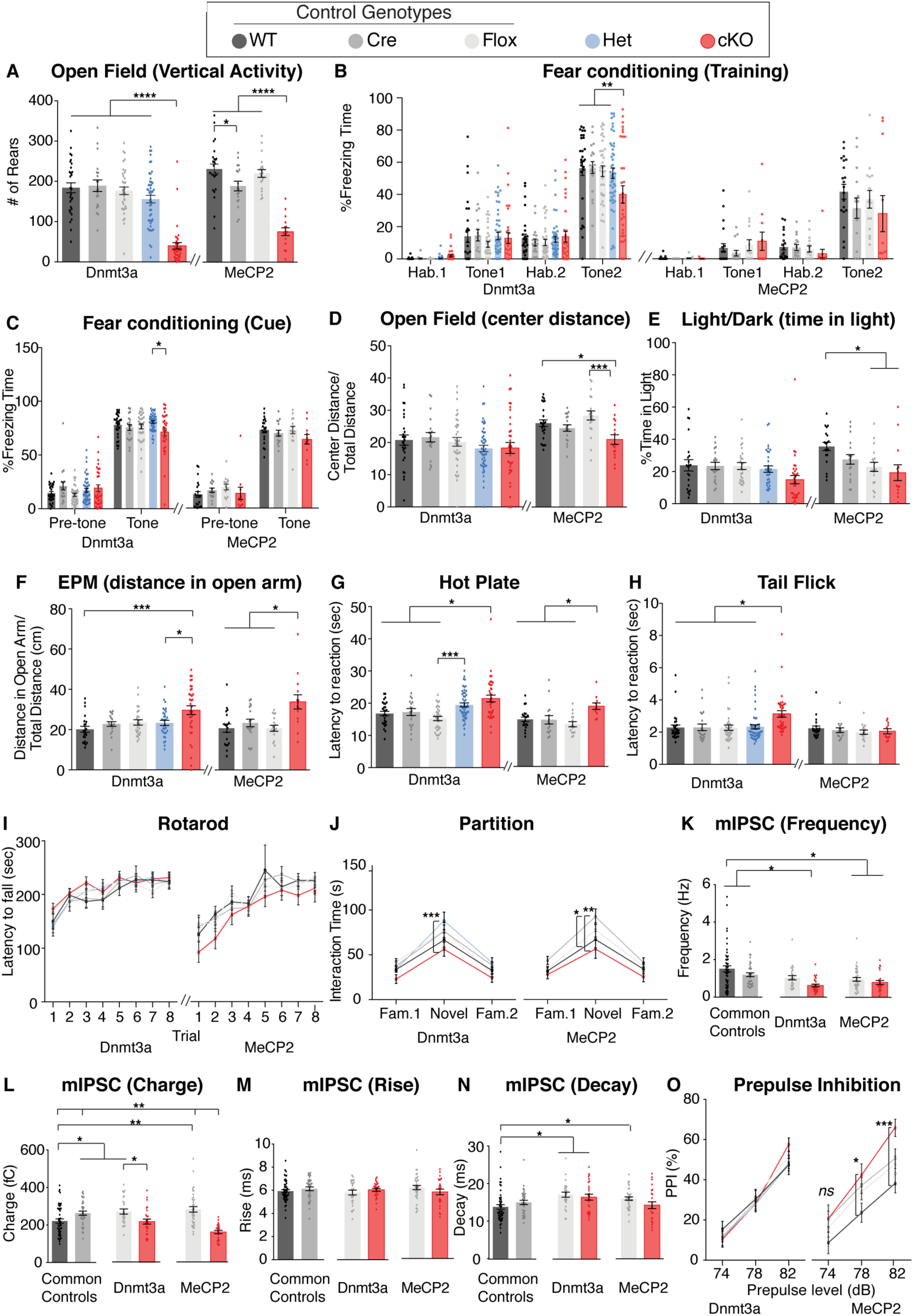
**A**) Both mouse lines showed decreased rearing in the Open Field test. **B)** Dnmt3a cKO mice showed impaired fear learning, whereas *Mecp2* cKO mice did not differ from control mice. **C)** Cue memory was normal in both cKO mice. **D-F)** Tests for anxiety-like behaviors (open field, light dark or elevated plus maze, respectively). **G)** Hot plate and **H)** tail flick testing for nociceptive pain in both cKO mice. **I)** Rotarod test for motor learning and coordination and **J)** the partition test for social interaction. **K-N**) Frequency, rise and decay measures from mIPSCs from the striatum. **O)** Only *Mecp2* cKO had a trend for increased pre-pulse inhibition. n= 11-50 per genotype (behavior), n= 5-9 mice per genotype with 24-50 neurons total (electrophysiology), *p<0.05, **p<0.01, ***p<0.001, ****p<0.0001. See Supplemental Table 1 for full numbers and statistics.

**Supplemental Figure 3.**
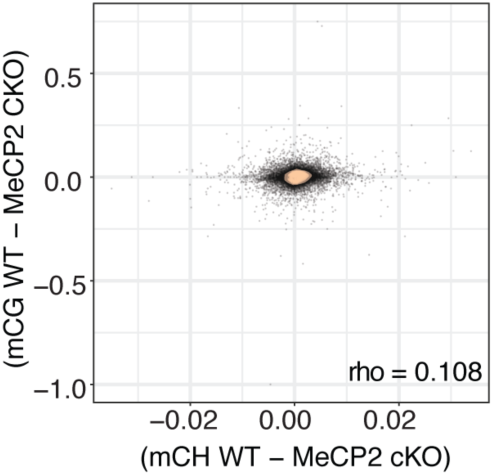
Plot of the difference in genome wide mCH versus mCG methylation between the WT and *MeCP2* cKO mice. Person correlation designated as rho. Consistent with stable methylation in the absence of MeCP2, the differences center at 0.

**Supplemental Figure 4.**
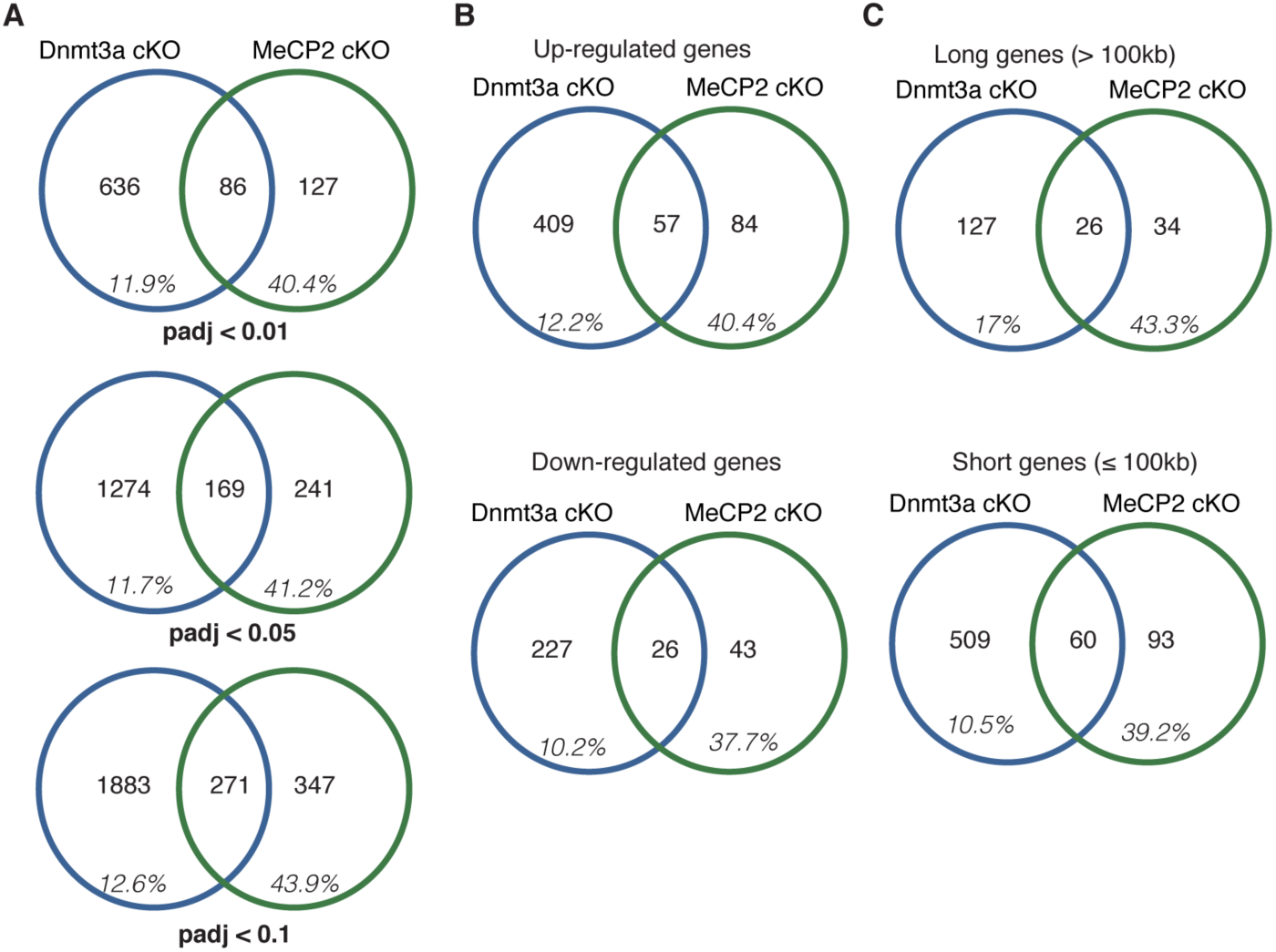
The percentage of DEGs that overlap between cKO mouse models broken down as a function of **A)** p-value, **B)** direction of change (down- or up-regulated), **C)** gene length.

**Supplemental Figure 5.**
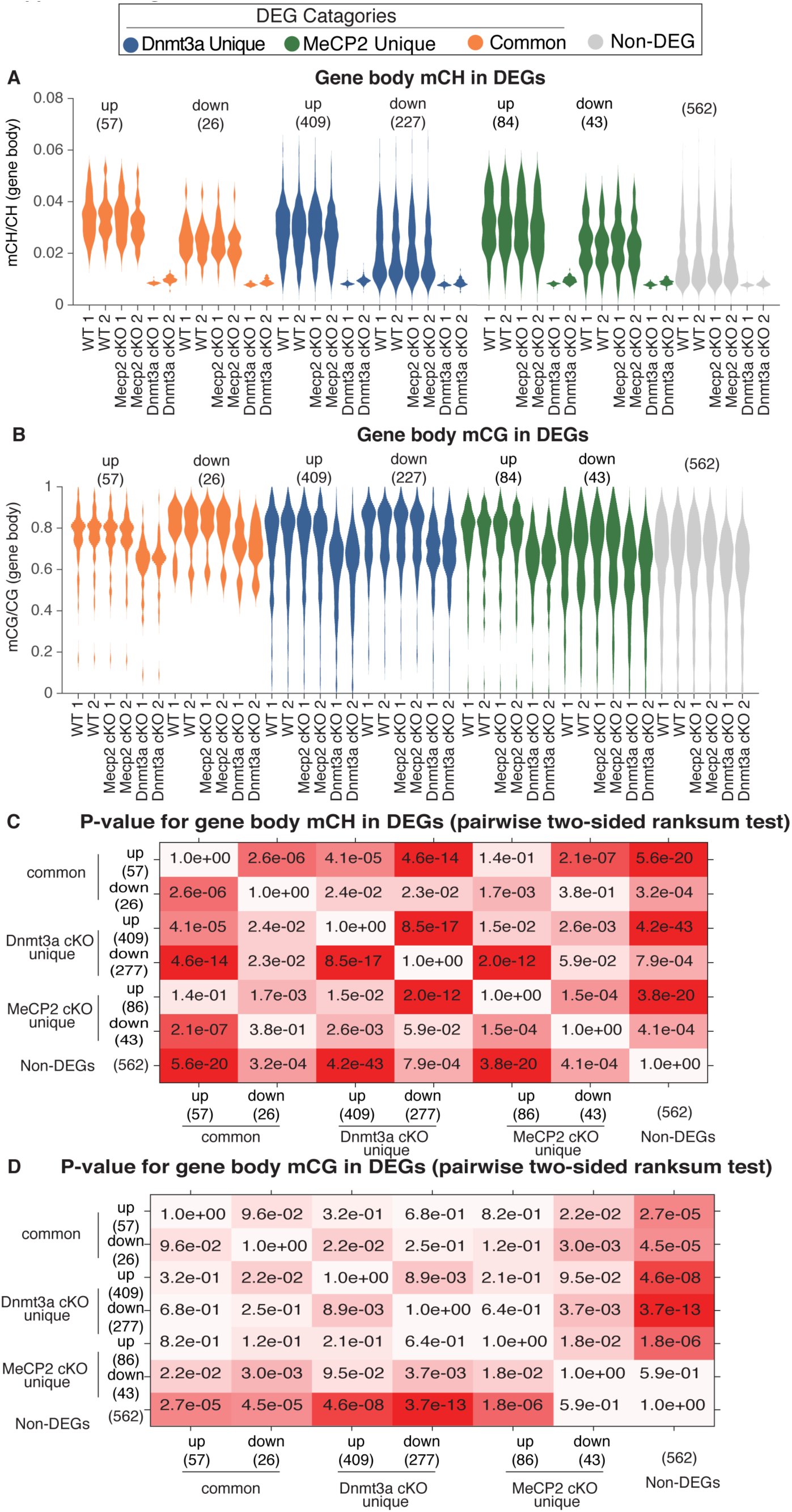
**A)** Gene-body mCH levels in different categories of DEGs in WT, *Dnmt3a* cKO and *Mecp2* cKO mice. **B)** Gene-body mCG levels in different categories of DEGs in WT, *Dnmt3a* cKO and *Mecp2* cKO mice. **C)** P-value matrix for comparison of mCH levels of different categories of genes illustrating significant differences versus non-DEGs. The boxes are colored such that more significant p-values are darker shades of red. **D)** P-value matrix for comparison of mCG levels of different categories of genes illustrating significant differences versus non-DEGs. As in C, the boxes are colored such that more significant p-values are darker shades of red.

**Supplemental Figure 6.**
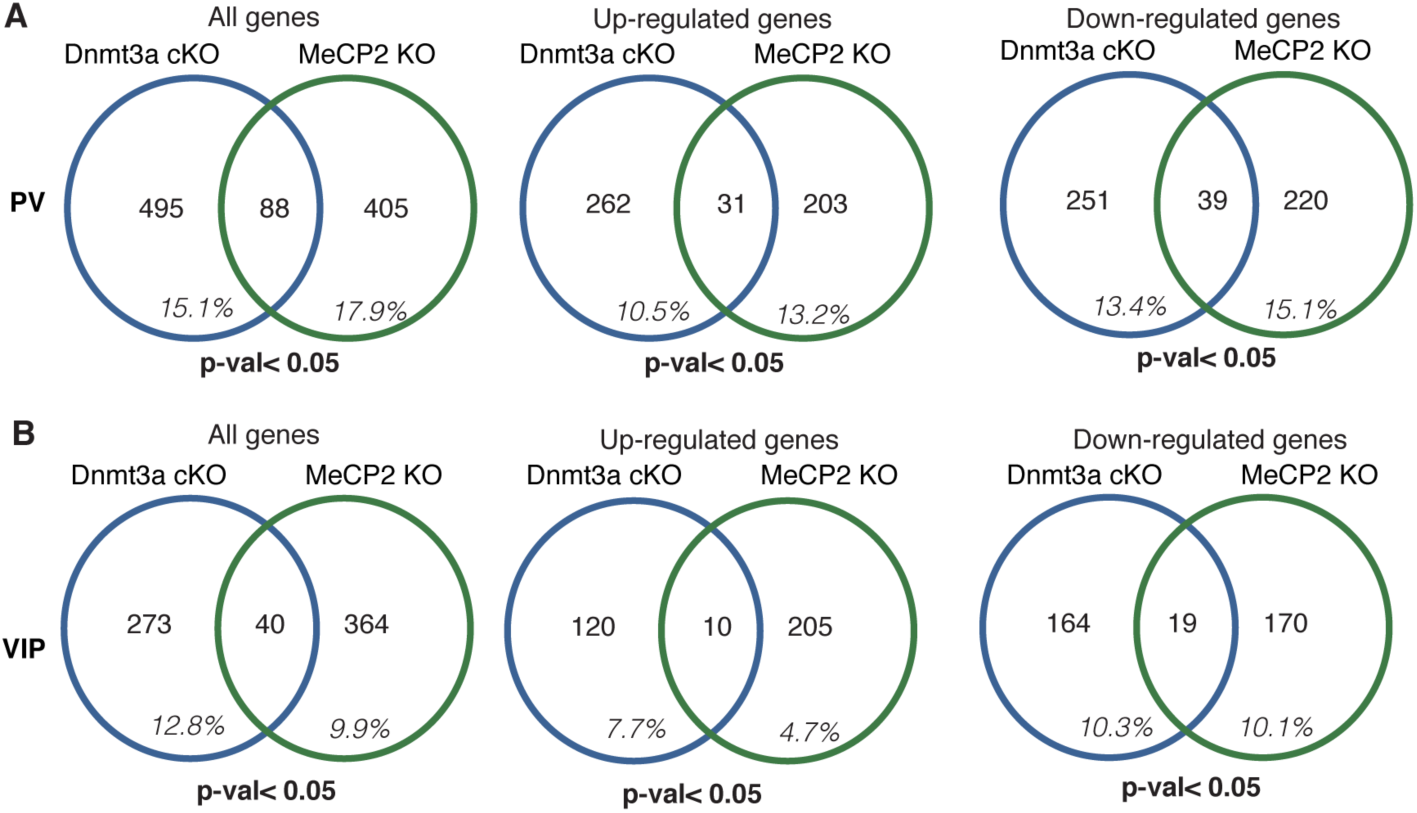
The percentage of DEGs that overlap between parvalbumin (PV) and vasoactive intestinal polypeptide (VIP) neurons in the mouse cortex from *Dnmt3a* cKO (Nestin-Cre) and *Mecp2* KO mouse models. The data are a re-analysis of single-nuclear RNA sequencing (19). The percentages are shown as all genes and genes broken down by direction of change (down- or up-regulated) for **A)** PV and VIP neurons. Consistent with our data there are few DEGs significantly misregulated in *Dnmt3a* cKO mice that are also significantly misregulated in the same neurons that lack MeCP2. Notably, our data show a significantly higher overlap of genes (40%) misregualted in *Mecp2* cKO that are also misregulated in the *Dnmt3a* cKO.

**Supplemental Table 1.** Numbers and statistics for all mouse behavioral assays and methylation datasets.

**Supplemental Table 2.** RNA-seq data and DEGs used in figures.

**Supplemental Movie 1.** Forepaw stereotypies are apparent in *Dnmt3a* cKO mice. Video shows an example of forepaw stereotypies in a *Dnmt3a* cKO male mouse at 8-week of age.

